# Blind source separation of event-related potentials using a recurrent neural network

**DOI:** 10.1101/2024.04.23.590794

**Authors:** Jamie A. O’Reilly, Hassapong Sunthornwiriya-Amon, Naradith Aparprasith, Pannapa Kittichalao, Pornnaphas Chairojwong, Thanabodee Klai-on, Edward W. Lannon

**Affiliations:** School of International & Interdisciplinary Engineering Programs, School of Engineering, King Mongkut’s Institute of Technology Ladkrabang, Bangkok 10520, Thailand; Department of Biomedical Engineering, School of Engineering, King Mongkut’s Institute of Technology Ladkrabang, Bangkok 10520, Thailand; Division of Pain Medicine, Department of Anesthesiology, Perioperative and Pain Medicine, Stanford University School of Medicine, 500 Pasteur Drive, Stanford, CA, United States of America

**Keywords:** Artificial neural networks, computational neuroscience, deep learning, electroencephalography (EEG), neural signal processing

## Abstract

Event-related potentials (ERPs) are a superposition of electric potential differences generated by neurophysiological activity associated with psychophysical events. Spatiotemporal dissociation of these signal sources can supplement conventional ERP analysis and improve source localization. However, results from established source separation methods applied to ERPs can be challenging to interpret. Hence, we have developed a recurrent neural network (RNN) method for blind source separation. The RNN transforms input step pulse signals representing events into corresponding ERP difference waveforms. Source waveforms are obtained from penultimate layer units and scalp maps are obtained from feed-forward output layer weights that project these source waveforms onto EEG electrode amplitudes. An interpretable, sparse source representation is achieved by incorporating L1 regularization of signals obtained from the penultimate layer of the network during training. This RNN method was applied to four ERP difference waveforms (MMN, N170, N400, P3) from the open-access ERP CORE database, and independent component analysis (ICA) was applied to the same data for comparison. The RNN decomposed these ERPs into eleven spatially and temporally separate sources that were less noisy, tended to be more ERP-specific, and were less similar to each other than ICA-derived sources. The RNN sources also had less ambiguity between source waveform amplitude, scalp potential polarity, and equivalent current dipole orientation than ICA sources. In conclusion, the proposed RNN blind source separation method can be effectively applied to grand-average ERP difference waves and holds promise for further development as a computational model of event-related neural signals.

## 1. Introduction

The main source of scalp electroencephalography (EEG) is considered to be synchronous post-synaptic potentials at dendrites of cortical pyramidal cells oriented perpendicular to the cortical surface [1], [2]. Electric fields due to synchronously active cortical patches of at least 6 cm become measurable from scalp electrodes [1]. Ongoing or “resting state” activity distributed broadly over the cerebral cortex makes raw EEG generally noisy and difficult to interpret. The event-related potential (ERP) technique circumvents this by averaging EEG signals recorded in response to repeated psychophysiological events in experiments designed to elicit associated neural signalling processes [3], [4]. This is made possible by the high temporal resolution of electromagnetic signal acquisition and the direct relationship between electric fields and underlying neurophysiology.

The ERP waveform recorded from an individual reflects part of their stereotyped neural response to a specific category of sensory-cognitive event. Brain activity associated with cognitive processes can be further isolated by subtracting ERPs from two or more event categories to produce ERP difference waves [5], [6]. These waveform subtractions are intended to remove features reflecting common sensory processes, while retaining features reflecting the cognitive processes of interest. ERPs and ERP difference waves from multiple subjects in a study are often averaged together to produce grand-average waveforms, and generalizations about brain activity are made from these grand-averages [7].

Conventional ERP waveform analysis involves measuring amplitudes and latencies. However, at any time instant the amplitude at a given scalp site can reflect the summation of multiple spatially separate cortical patches with temporally overlapping activity [3]. Identification of these underlying source waveforms and their corresponding scalp distributions is desirable for interpreting ERPs and enhancing conventional waveform analysis. When no prior information is given about the processes involved in generating the observed signal of interest (i.e., the ERP waveform) the procedure of identifying underlying source waveforms is referred to as “blind source separation” [8]–[10].

The blind source separation method most widely used in EEG analysis is independent component analysis (ICA), which deconstructs independent EEG signals into an equal number of statistically independent components (ICs) [10]–[12]. ICA optimizes an unmixing matrix to transform EEG signals into IC signals with minimal shared information. The inverse mixing matrix, referred to as the unmixing matrix, transforms ICs into EEG signals, thus providing the weights (spatial filter or “scalp map”) required to compute scalp projections from each individual IC source waveform [12]. This technique is particularly effective for preprocessing EEG by correcting artifacts such as eye-blinks [10], [13], [14], but can also identify separable brain processes for further examination [9], [11], [12]. Blind source separation of grand-average ERP waveforms can be achieved by applying ICA to a database of ERPs from different subjects, which can be referred to as group-ICA [15], [16]. Doing so produces source waveforms and scalp maps that generalize across subjects. However, two limitations of ICA pertinent to this study are: (i) the number of sources is fixed by the requirement for square mixing/unmixing matrices, and (ii) the polarity of source waveforms and scalp maps can be inverted, obscuring direct visual analysis [12], [17], [18].

A recurrent neural network (RNN) can overcome these limitations of ICA for ERP data by (i) allowing more sources than the number of independent EEG channels, which can be optimized for sparsity with L1-norm regularization, and (ii) using a rectifying activation function to enforce positive amplitudes from source waveforms. RNNs for analysing ERP waveforms have recently been developed for modelling auditory evoked potentials from mice [19], [20], human ERPs [21], [22], and combining with a convolutional neural network (CNN) to study visual ERPs [23]. RNNs can also be used for distributed source reconstruction from MEG [24], EEG [25], and simultaneously recorded MEG-EEG [26]. These previous studies demonstrate some of the ways that RNNs can be used for analysing event-related neural signals. The current paper extends this work by developing a new method for separating ERP source waveforms and their corresponding scalp maps. This new method is applied to a collection of ERP difference waveforms from multiple subjects, thus generalizing grand-average data. However, this method could be adapted to model single-subject ERP data. ICA blind source separation was applied to the same data and the results from both methods were compared.

## 2. Materials and Methods

### 2.1. Event-related potential data

Data from the ERP Compendium of Open Resources (ERP CORE) were analysed in this study [27]. This database includes normative data recorded from 40 healthy adult subjects for some of the most well-established ERP paradigms. Twenty-eight EEG electrode recordings were used, named by the International 10-20 System: FP1, FP2, F3, Fz, F4, F7, F8, FC3, FCz, FC4, C5, C3, Cz, C4, C6, CPz, P7, P3, Pz, P4, P8, PO7, PO3, PO4, PO8, O1, Oz, and O2. Some artifacts in continuous EEG were corrected with ICA [13] and ICLabel [28] by automatically removing components classified as “muscle artifact”, “eye blink”, “heart beat” or “channel noise”. Corrected signals were then band-pass filtered from 0.1 to 20 Hz and re-sampled to 100 Hz before re-referencing to the channel average. Relevant epochs were extracted from 0.2 s before stimulus onset to 0.8 s after stimulus onset. Resulting epochs were averaged to produce the ERP from each subject. Some subjects were removed due to excessive artifacts following recommendations provided in ERP CORE.

From the passive auditory oddball paradigm, 80 dB standard ERPs were subtracted from 70 dB deviant ERPs to produce mismatch negativity (MMN) difference waveforms from 39 subjects. From the face perception paradigm, car ERPs were subtracted from face ERPs to obtain face N170 difference waveforms from 37 subjects. From the word pair judgement paradigm, related-word ERPs were subtracted from unrelated-word ERPs to produce N400 difference waveforms from 39 subjects. From the active visual oddball paradigm, frequent non-target ERPs were subtracted from rare target ERPs to produce P3 difference waveforms from 34 subjects. Altogether the resulting dataset included 149 ERP difference waves.

### 2.2. Recurrent neural network modelling

The ERP difference waveforms were scaled by 10^6^ and combined to form a label tensor 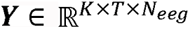 with *K* = 149 difference waveforms, *T* = 101 time-samples, and *N_eeg_* =28 EEG channels. The label tensor ***Y*** may be considered as a set of difference waveforms {*Y*_1_, *Y*_2_, …, *Y_K_*} where 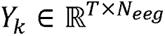 is a difference waveform obtained from an individual subject, and 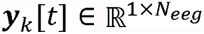 is a row vector containing amplitudes from a single time-sample of an individual ERP difference waveform. A matching input tensor 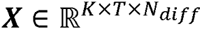 was constructed with *N_diff_* =4 channels containing step-pulse signals representing each type of difference waveform. Similarly, ***X*** is composed of {*X*_1_, *X*_2_, …, *X_K_*} where 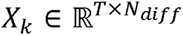 and 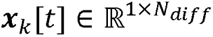. Only one of the four channels in ***X*** had a unit step pulse from 0 to 0.2 ms during each of 149 instances of ***Y*** (first channel: MMN, second channel: N170, third channel: N400, and fourth channel: P3); the other three channels not associated with the respective ERP difference waveform were zero. These data are illustrated on Figure 1 below a block diagram of the RNN, which learned a function *f*(***X***) = ***Ŷ***, where the mean-squared error (MSE) between ***Y*** and ***Ŷ*** was minimized by gradient descent.

**Figure 1.**
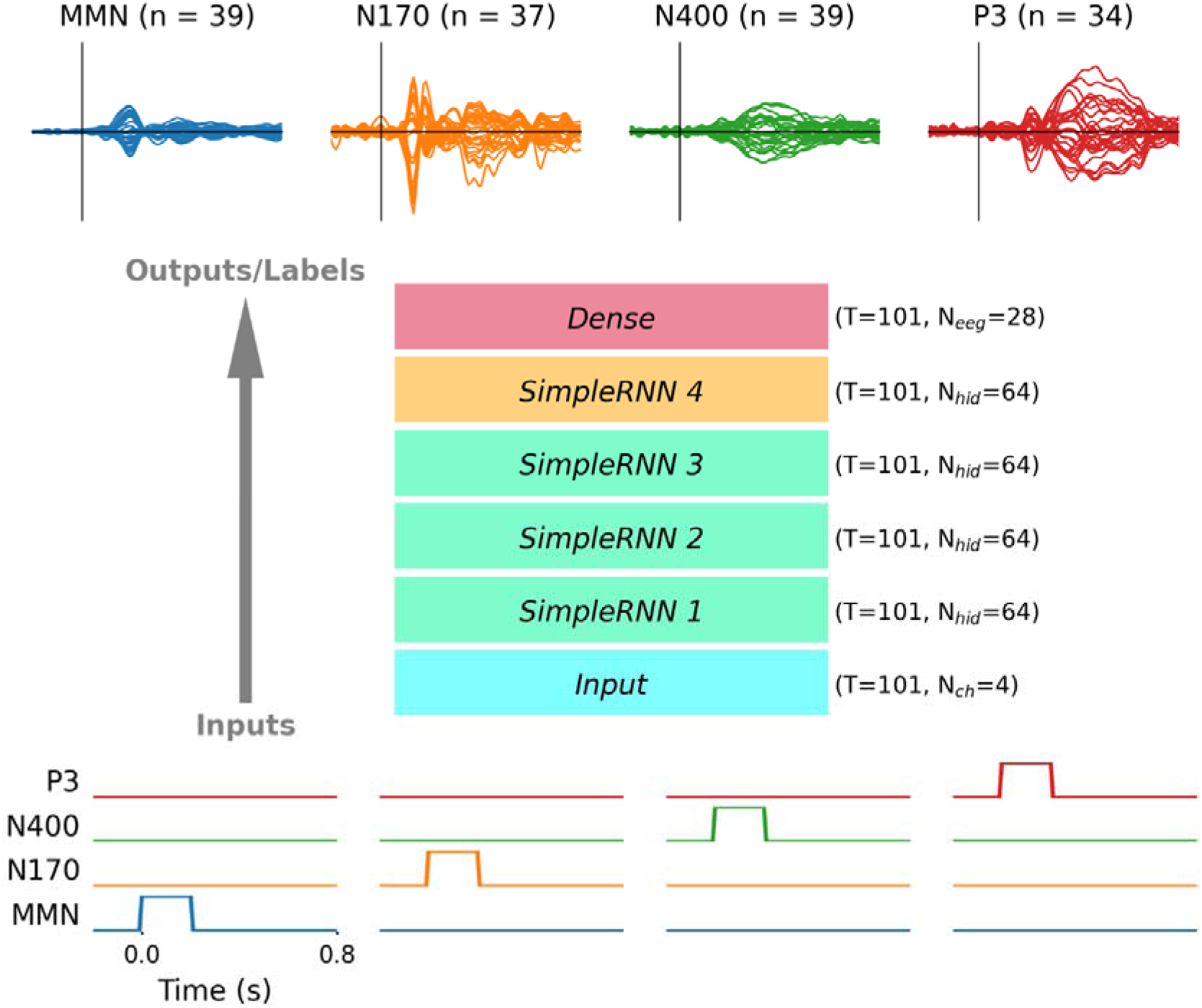
Diagram illustrating the RNN modelling approach. Four model inputs (bottom) were paired with ERP difference waveforms from 28 EEG channels used as labels (top) to train the RNN by supervised learning. From left to right, inputs represent MMN, N170, N400, and P3 difference waves with a unit step pulse high from 0 to 0.2 s on a specific channel. The model architecture consisted of an input layer, followed by four recurrent layers, and a fully-connected output layer. The temporal structure of T = 101 time-samples was preserved throughout the network, while the number of channels at different layers varied from N_ch_ = 4 at the input, to N_hid_ = 64 for hidden layers, and N_eeg_ = 28 at the output. A one-to-many mapping between each input representations and multiple ERP difference waveforms caused model outputs to approximate the grand-average ERP difference waves. Source signals were obtained from the *SimpleRNN 4* layer, and weights learned by the *Dense* layer project these source waveforms onto scalp EEG channels. Difference waves plotted at the top for illustrative purposes are grand-averages from subjects, with 28 channel ERPs plotted from −0.2 to 0.8 s (the same time scale as input signals shown at the bottom).

The RNN architecture consisted of an *Input* layer, four *SimpleRNN* hidden layers with *N_hid_* =64 hidden units each, and a *Dense* output layer with *N_eeg_* units (layer names in italics reflect TensorFlow nomenclature). The *Input* layer configures the network to receive input data with *T* time-samples and *N_diff_* channels. The first *SimpleRNN* layer implements equation (1), where 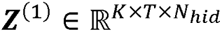 is the output tensor, 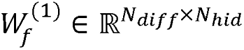 are forward weights, 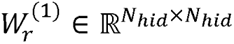 are recurrent weights, and 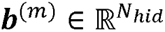 is the bias vector. The second and third *SimpleRNN* layers implement equation (2), which is identical to (1) but takes input from the preceding layer, so 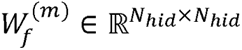 for hidden layer *m* = 2,3,4. The fourth *SimpleRNN* layer implements equation (3), which is essentially the same as (1) and (2) but without the bias vector. The *Dense* feed-forward output layer also excluded a bias vector, implementing equation (4) where 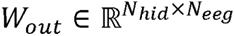. Each row of coefficients in *W_out_* defines the contribution of each source signal in ***Z***^(4)^ to each electrode channel in ***Y***, effectively acting as a spatial filter or scalp map. The rectified linear unit *ReLU*(·) activation function returns zero when its argument is negative and returns the argument when its argument is positive, enforcing output tensors to be non-negative.

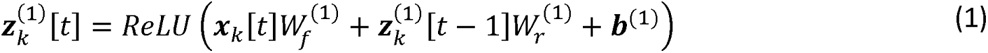

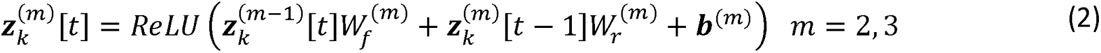

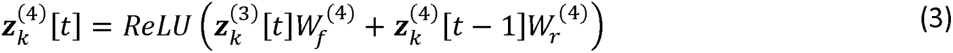

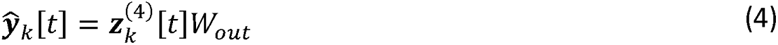

Model training was completed in two phases. In the first phase, the RNN was trained solely to minimize MSE loss in equation (5), where *K* = 149 was the number of difference waveforms and *T* = 101 was the number of time samples. In the second phase, the pre-trained RNN from phase one was fine-tuned with an L1-norm penalty (*activity_regularizer* coefficient, *α_L_*_1_ = 0.0001) applied to ***Z***^(4)^ and an L2-norm penalty (*kernel_regularizer* coefficient, *α_L_*_2_ = 0.01) applied to the output layer weights, as in equation (6). This made ***Z***^(4)^ sparse and constrained the range of *W_out_*. The adaptive-moment estimation (Adam) optimizer was used with default hyperparameters (*learning_rate* = 0.001, *beta_1* = 0.9, and *beta_2* = 0.999). Batch size was *K*, and the RNN was trained for 5000 iterations or until 250 epochs had elapsed without reduction in loss. After completing this training procedure, source waveforms were obtained from units with non-zero elements in ***Z***^(4)^ and scalp maps were obtained from their associated rows in *W_out_*.

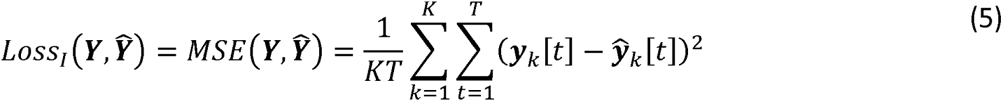

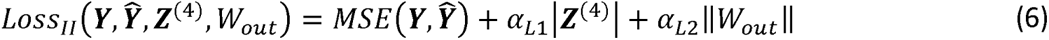

### 2.3. Independent component analysis decomposition

Independent component analysis was performed to extract IC sources from the same dataset of ERP difference waves used with the RNN method. The data in ***Y*** were structured as an *EpochsArray* object using MNE-Python [29] and the *extended infomax* ICA algorithm [30], [31] implemented in MNE-Python was applied to this object. This approach can be referred to as group ICA because the data used to derive the ICA decomposition came from a group of subjects [16]. The data matrix rank was 27 because EEG channels had common-average referencing. The principal component with least explained variance was removed to avoid rank deficiency before applying ICA, thereby preventing “ghost components” [17], [32]. The subsequent full-rank ICA decomposition yielded 27 ICs that were averaged across subjects for each ERP difference wave [i.e., MMN (n = 39), N170 (n = 37), N400 (n = 39), and P3 (n = 34)] and multiplied by their associated scalp maps to reconstruct grand-average difference waves (see Figure 3).

### 2.4. Data analyses

Pearson’s correlation coefficient (r) and MSE were used to compare waveforms reconstructed by RNN and ICA sources with grand-average ERP difference waveforms. Correlation was also calculated between source signals and scalp topographies, as described below. The *scipy.stats.pearsonr* and *numpy.mean* functions were used to calculate r and MSE, respectively.

Shannon entropy (SE) of each source projection’s correlation with four difference waveforms was used to quantify the degree of component-specificity of each source; SE of zero indicates that the source was associated with only one ERP difference wave, and higher values indicate correlations across multiple difference waves. The *scipy.stats.entropy* function was used to compute SE. For any ICA-derived sources that had negative correlation with some difference waveforms, their correlations were transformed into absolute values before calculating SE.

Mutual information (MI) between each pair of RNN and ICA sources was also computed. MI is equal to zero only if the two signals are independent, and higher values indicate greater dependency between the pair of signals. Source signals associated with four difference waves were compared at once by first concatenating them. The *sklearn.feature_selection.mutual_info_regression* function was used to compute MI.

A similarity score (SS) was calculated for each pair of RNN and ICA sources using equation (7). Here, ***x****_i_* ∈ ℝ^404^ was the row vector containing concatenated signals from source *i*, associated with four difference waveforms [i.e., 404 = *T* x *N_diff_*], ***x****_j_* ∈ ℝ^404^ contained equivalent signals from source *j*, 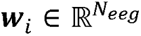 was the scalp map for source *i*, 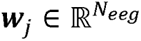 was the scalp map for source *j*, *MI*(·) was the mutual information function, and *r*(·) was Pearson’s correlation function. Taking the absolute values of correlation between pairs of ***x*** and ***w*** vectors accounted for source signals and topographic maps produced by RNN and ICA methods potentially being anticorrelated; this can happen because ICA allows biphasic signals [12].

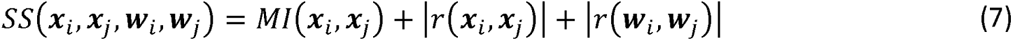

### 2.5. Software

Python 3 with matplotlib 3.5.2 [33], mne 1.4.0 [29], numpy 1.25.2 [34], scikit-learn 1.3.2 [35], scipy 1.11.2 [36], and tensorflow 2.14.0 [37] were used in this study. *Data and code developed in this study will be shared in a public repository at publication*.

## 3. Results

RNN performance after each training phase is reported in Table 1. In each training phase 5000 iterations were completed. After the first training phase, RNN outputs correlated almost perfectly with ERP difference waveforms, which all had r > 0.98 and MSE < 0.005. This fit was achieved using 64 source waveforms in the penultimate layer. After the second training phase (i.e., fine-tuning the network with L1-norm regularization), only 11 source signals were required to fit the training data, and the other 53 hidden units in the *SimpleRNN* 4 layer had zero amplitude. Evaluation metrics comparing model outputs with grand-average ERP difference waves deteriorated slightly while remaining decent; all r > 0.9 and MSE < 0.035. Performance of ICA source reconstruction of ERP difference waveforms is also reported in Table 1. These are also highly correlated (all r > 0.96), but have slightly higher MSE (all < 0.066) than RNN source reconstructions.

**Table 1.** Summary of RNN performance after two training phases and ICA performance.

| Training phase<sup>1</sup> | Iterations | Nonzero sources | MMN |  | N170 |  | N400 |  | P3 |  |
| --- | --- | --- | --- | --- | --- | --- | --- | --- | --- | --- |
|  |  |  | r | MSE | R | MSE | r | MSE | r | MSE |
| 1 | 5000 | 64 | 0.989 | 0.00172 | 0.995 | 0.00486 | 0.994 | 0.0024 | 0.997 | 0.00416 |
| 2 | 5000 | 11 | 0.907 | 0.0147 | 0.967 | 0.0345 | 0.918 | 0.0344 | 0.983 | 0.0274 |
| ICA<sup>2</sup> | <500 | 27 | 0.965 | 0.0119 | 0.994 | 0.0651 | 0.988 | 0.0246 | 0.997 | 0.106 |

<sup>1</sup>In training phase 1 equation (5) was minimized; in training phase 2 equation (6) was minimized.
<sup>2</sup>*Extended Infomax* ICA algorithm implemented in MNE-Python.

When referring to RNN sources the remaining parts of the results, and discussion and conclusions sections implicitly refer to those obtained after the second training phase, unless otherwise stated. RNN sources, associated scalp maps, and reconstructed difference waveforms for channels of interest are plotted in Figure 2. The ICA method produced 27 independent sources, whose time-courses, scalp maps, and reconstructed ERP difference waves for channels of interest are plotted in Figure 3. Reconstructed ERPs for all channels from RNN sources are plotted in Figure S1 and those from ICA sources are plotted in Figure S2.

**Figure 2.**
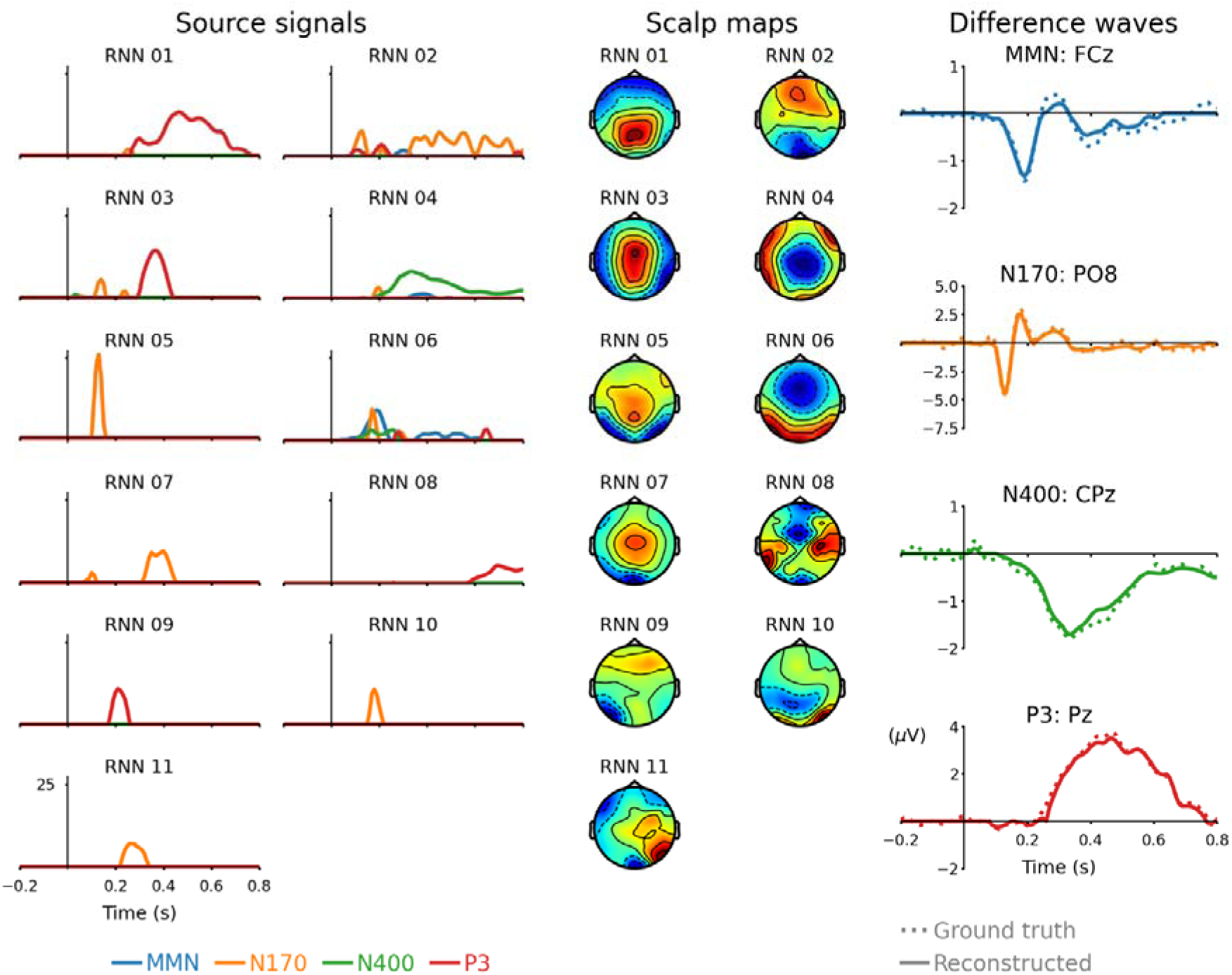
Source signals (left), scalp maps (middle), and reconstructed ERP difference waveforms (right) from RNN sources. Sources are arranged in reverse order of overall correlation between source projections and ERP difference waveforms. Scalp maps are each scaled independently and symmetrically about zero for maximum positive (dark red) and maximum negative (dark blue) values. Dashed lines represent ground truth difference waveforms; solid lines represent reconstructed difference waveforms from the projections of these 11 RNN sources.

**Figure 3.**
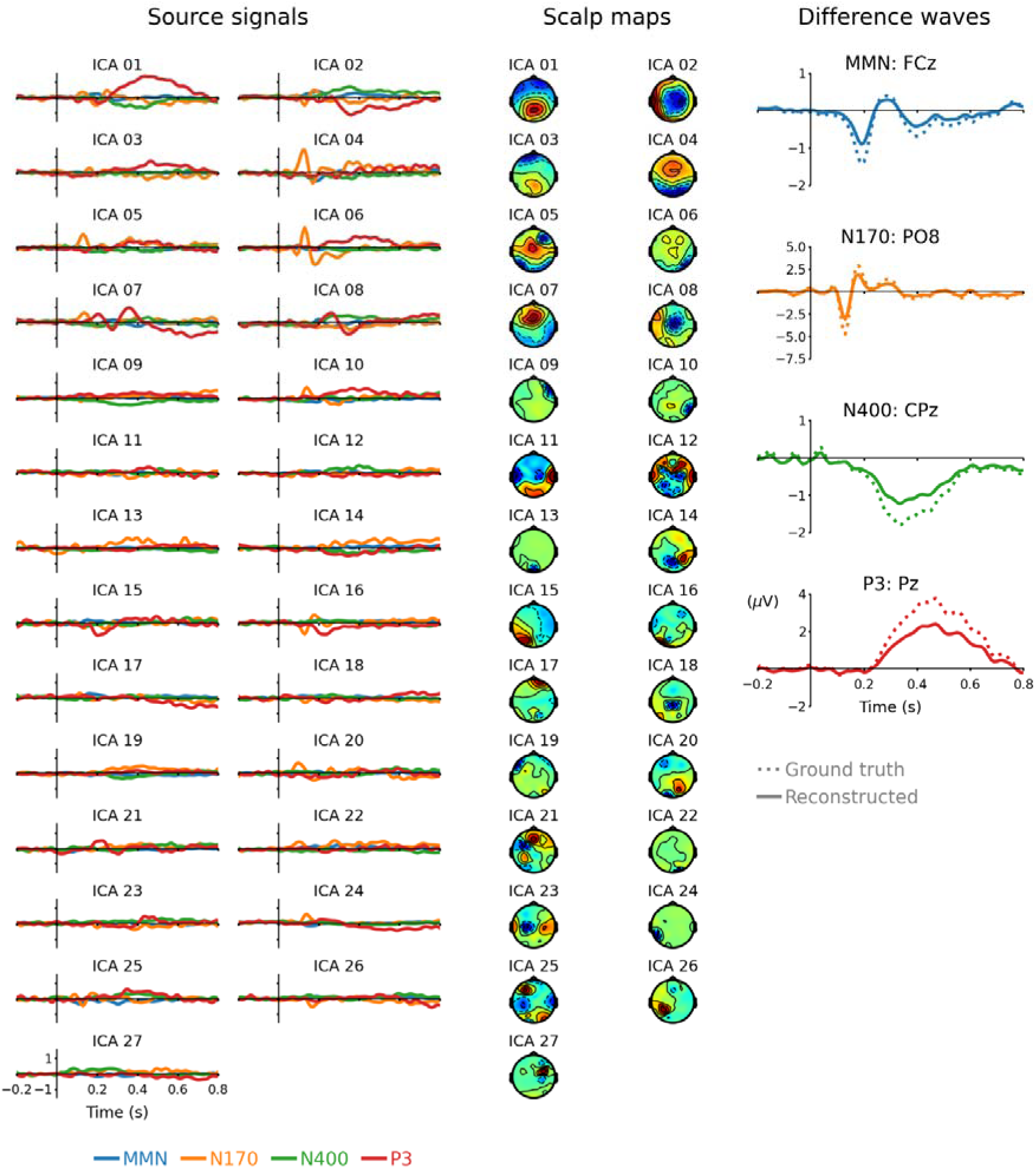
Source signals (left), scalp maps (middle), and reconstructed ERP difference waveforms (right) from ICA blind source separation. Difference waveforms reconstructed by ICA sources have slightly lower amplitudes than the ground truth; this may be because source signals for each subject were averaged before reconstructing these grand-average ERP difference waveforms. Separate figures for each RNN and ICA source showing source waveforms, scalp maps, and correlation of projections with four ERP difference waves can be viewed from: *link will be shared with published version*.

Correlation and MSE between each source’s projection and ERP difference waveforms are given in Table S1, which also reports SE and the most similar source for each source. It can be seen that RNN sources generally have lower entropy (μ = 0.36, σ = 0.43) than ICA sources (from 27 sources: μ = 1.2, σ = 0.138). Mutual information was also lower for RNN sources (from 55 pairwise comparisons: μ = 0.0174, σ = 0.0275) than ICA sources (from 351 pairwise comparisons: μ = 0.3, σ = 0.123). Pairwise comparisons between all sources in terms of MI between signals are represented in Figure S3, r between signals are represented in Figure S4, and r between scalp maps are represented in Figure S5.

Computed similarity scores for each pair of RNN and ICA sources are displayed in Figure 4. Three pairs of RNN and ICA sources are reciprocally most similar with each other based on this analysis. These are RNN 01 and ICA 01, whose projections both correlate strongly with P3; RNN 02 and ICA 04, whose projections both correlate well with N170 (although ICA 04 projections also correlate well with MMN); and RNN 04 and ICA 02, whose projections both correlate highly with N400 (although ICA 02 projections also correlate well with other ERPs). These correspondences indicate that both methods extract some similar features from the data. Notable similarity was also seen from RNN and ICA sources extracted from the same data without correcting eye-blink artifacts, examples of which are plotted in Figure S6; here, ICA 01 captured both phases of the eye-blink source, RNN 01 captured negative scalp potential phases of the eye-blink, and RNN 02 captured positive scalp potential phases of the eye-blink.

**Figure 4.**
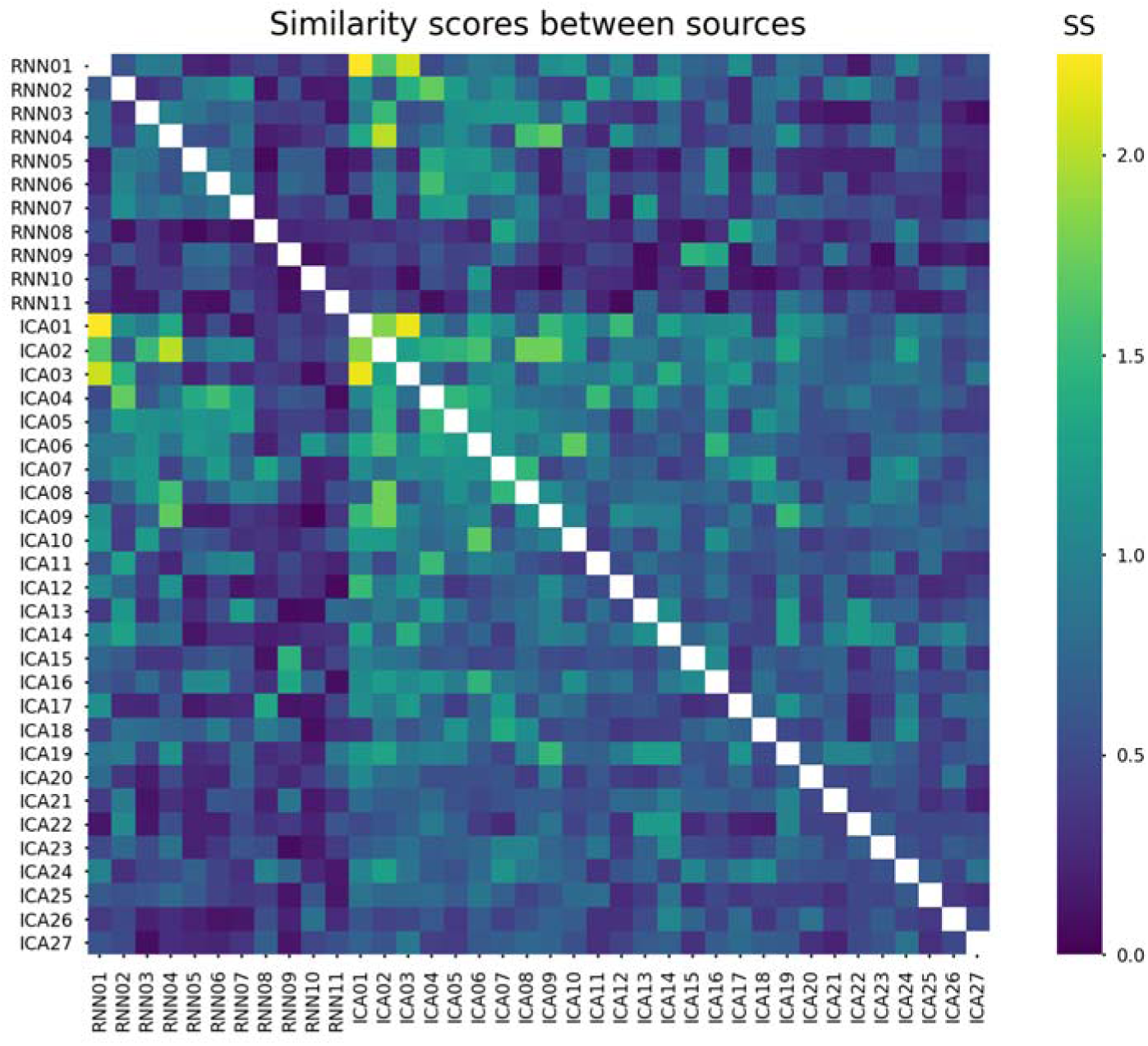
Similarity scores between each pair of sources. These SS values were calculated using equation (7) and reflect contributions from MI (Figure S3) and r (Figure S4) between source waveforms and r between scalp maps (Figure S5). Generally, SS is lower within RNN sources than within ICA sources, and higher between RNN and ICA sources than within RNN sources. For the source in each row, its corresponding most similar source was determined from the column with the highest SS. Diagonal elements are omitted.

RNN hidden unit activations from all layers and outputs associated with each ERP difference wave are plotted in Figure 5. Equivalent data from the RNN after the first training phase are plotted in Figure S7. The RNN model transforms simple input step signals into output waveforms corresponding to grand-average ERP difference waves from 28 EEG channels. The first hidden layer (*SimpleRNN* 1) outputs depict ongoing oscillations that are perturbed when its input signal changes. The last hidden layer (*SimpleRNN* 4) outputs plotted in Figure 5 are the source waveforms in Figure 2 grouped by difference wave type. Three hidden units were active for MMN (02, 04, and 06); nine were active for N170 (01, 02, 03, 04, 05, 06, 07, 10, and 11); three were active for N400 (03, 04, and 06); and six were active for P3 (01, 02, 03, 06, 08, and 09). Mathematical relationships among these source signals can be studied using the weights of the trained RNN. For example, the recurrent weights from the last hidden layer (*W*_r_^(4)^, represented in Figure S8) can be interpreted as functional connectivity between source signals, and signals from the preceding layer (*SimpleRNN* 3) can be viewed as drivers of each source waveform.

**Figure 5.**
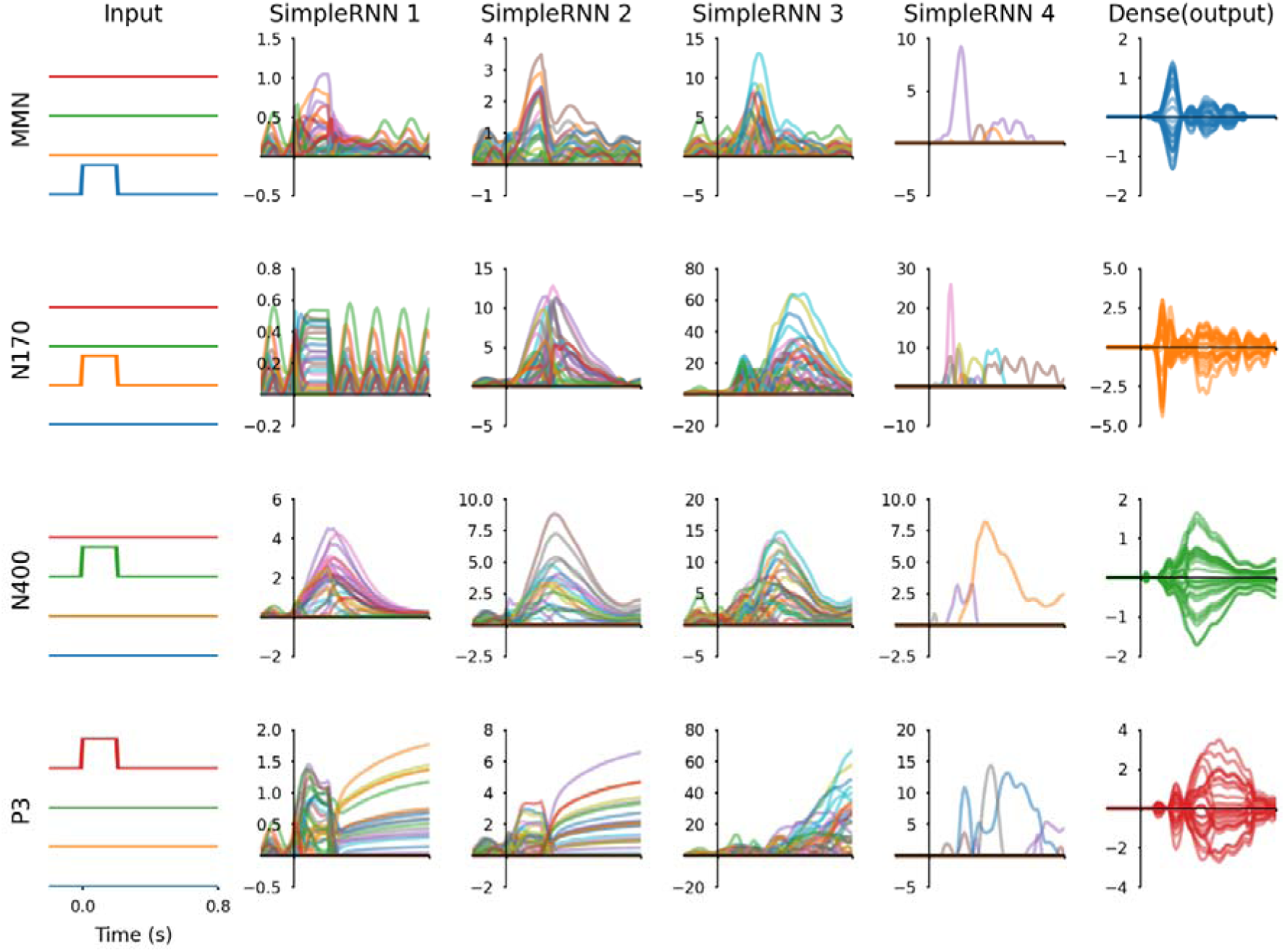
Inputs, hidden unit activations, and output signals from the RNN. The rows show model responses for four difference waveforms, the columns show all of the signals at each layer of the RNN. Outputs effectively reproduce ERP difference waves from 28 EEG channels; these are colour-coded to match the associated input channel, although hidden unit activity traces are coloured arbitrarily. *SimpleRNN* 4 signals active for MMN are 02, 04, and 06; for N170 are 01, 02, 03, 04, 05, 06, 07, 10, and 11; for N400 are 03, 04, and 06; and for P3 are 01, 02, 03, 06, 08, 09. All signals are plotted with the same time range. Input signals are binary, RNN hidden unit signals are dimensionless, and RNN outputs correspond with ERP amplitudes in microvolts. A corresponding figure for the RNN after the first training phase is presented in Figure S8.

## 4. Discussion

### 4.1. Theoretical validity of RNN sources

Interpreting results from blind source separation of grand-average data from multiple paradigms presents a dilemma: whether sources that contribute to multiple ERPs reflect shared neurophysiological/cognitive operations or distinct operations with shared scalp distributions. Both RNN and ICA methods produced sources involved in reconstructing multiple ERP difference waveforms, although RNN sources tended to be more specific. Sensory modality and theories from cognitive neuroscience can be considered to rationalize these findings. However, justifying all of the sources involved is particularly challenging for ICA, which had 26 (MMN), 27 (N170), 25 (N400), and 25 (P3) source projections that were positively correlated with multiple ERP difference waves. Deciphering these source signals is problematic, and some are possibly artifacts of ICA. In contrast, the sparser representation of RNN sources is simpler and easier to describe, hence more likely to be useful for generalizing these data. Summaries of RNN source findings for each ERP difference waveform are given below, before directly comparing RNN and ICA methods.

#### MMN

The RNN required three non-zero sources to produce the MMN difference waveform. These are labelled 02 (r = 0.175), 04 (r = 0.269), and 06 (r = 0.87) in Figure 2. Source 06 can be considered the main source of interest for MMN. These sources also had relatively high entropy compared with other RNN sources (SE = 0.843, 0.822, and 1.124, respectively), which suggests that the sources underlying MMN may also play a role in N170, N400, and P3. Functional overlap between the sources of these four ERP difference waveforms is plausible, as each of these ERPs can be interpreted in the context of predictive processing [38] and conscious perception [39]. However, it is also possible that functionally distinct but spatially indistinguishable (using 28 EEG channels) sources are being represented by different source waveforms with shared scalp distributions.

#### N170

The RNN required nine sources to reconstruct N170: 01, 02, 03, 04, 05, 06, 07, 10, and 11. Of these, projections from source 02 had the highest correlation (r = 0.746) with the N170 waveform; four sources (05, 07, 10, and 11) were only active for N170 (SE = 0). Several of these signals overlap around the time range of 0.17 s post-stimulus, but source 05 stands out as the most likely to coincide with the negative peak from the bilateral fusiform face area (FFA) prominently observed from channels PO7 and PO8 [40], [41]. Source 10 has a scalp distribution consistent with the positive deflection observed at approximately 0.18 s from the FFA. The temporal dependency between sources 05 and 10 is illustrated by recurrent weights in Figure S8, which show that source 05 suppresses activity of source 10, whereas source 05 is comparatively unaffected by source 10; indicating unilateral influence following the temporal sequence of these source waveforms, with the earlier source affecting the later source.

#### N400

Three RNN sources generated N400: 03 (r = 0.042), 04 (r = 0.889), and 06 (r = 0.301). Source 04 peaks at 0.33 s then steadily decays. It has a left-dominant scalp distribution, consistent with prior literature on N400 [42], [43]. Source 06 peaks at 0.26 s and has a posterior scalp distribution, suggesting differential processing in visual association areas preceding activation of source 04. Recurrent weights displayed in Figure S8 show that sources 06 and 04 tend to suppress each other, which partly explains why source 04’s amplitude increases as source 06’s amplitude decreases. This temporal dependency between sources, and their corresponding scalp maps, suggests that the N400 ERP difference is underpinned by two separate components.

#### P3

Five RNN sources had projections that correlated with P3: 01 (r = 0.868), 02 (r = 0.166), 03 (r = 0.509), 06 (r = 0.112), 08 (r = 0.205), and 09 (r = 0.2). Sources 03 and 01 are candidates for P3a and P3b subcomponents, respectively, whose time courses and scalp maps agree strikingly with illustrations in [44]. Source 03 peaks at 0.37s and has a slightly more anterior positive scalp distribution than source 01, which peaks at 0.46 s and has slightly posterior positive scalp distribution. Although activity of these sources overlaps and their scalp distributions are similar, the RNN cleanly distinguishes them. The other sources (02, 06, 08, and 09) may reflect neural processes involved in producing P3 difference waveforms that are yet to be fully characterised.

### 4.2. Comparisons between RNN and ICA methods

#### 4.2.1. General comparison

The RNN method effectively deconstructs ERP difference waveforms into a collection of underlying source waveform and corresponding scalp maps, producing results comparable with ICA. Weights learned by the feed-forward output layer of the RNN do not change, making the corresponding scalp maps for each source waveform static for the duration of the ERP. This matches the assumption of ICA that independent sources of EEG are physically static throughout the data [12], [17].

The EEG inverse problem is generally assumed to be underdetermined, with there being many more neural sources than recording electrodes [45], [46]. As seen from Figure S8, the RNN method can produce more sources than channels for reconstructing ERP signals when trained without L1-norm regularization of source signals included with the loss function. However, the number of sources decreased from 64 to 11 by including L1-norm regularization, which attenuates source waveforms that are less consequential for fitting the data. In contrast, the number of ICs produced by ICA is fixed to equal the number of independent channels (i.e. *N_eeg_* 1=27 for 28 EEG channels with common average reference). This feature of ICA may produce excessive sources for ERP data, some of which can be artifacts. RNN performance metrics deteriorate after the second training phase compared with the first phase or ICA performance (Table 1), reflecting a trade-off between source representation complexity and these performance metrics. ERP reconstructions from more complex source representations achieve higher correlation and lower MSE when evaluated against the grand-average ERP difference waves, although this partially reflects fitting noise (e.g., in the baseline window). Therefore, a marginal performance deterioration may be considered an acceptable compromise for a sparser, more interpretable source representation.

RNN source signals were qualitatively smoother than ICA sources. By only having positive amplitudes they also had less ambiguity between source waveform and scalp projection polarity. Fewer, cleaner source signals, with directly proportional relationships with scalp potential distribution, make RNN sources generally easier to inspect and interpret. Nevertheless, sources from both methods reconstructed ERP difference waves that correlated highly with ground-truth grand-average ERP difference waves. However, despite having comparable correlation with ground-truth ERPs, waveforms reconstructed by ICA sources had higher MSE than those reconstructed by RNN sources. This could be due to ICA source signals being averaged across subjects, to obtain grand-average source waveforms, before reconstructing grand-average difference waves; rather than reconstructing individual subject ERPs then averaging them to obtain the grand-average ERP.

#### 4.2.2. Source waveform polarity

The RNN enforced source waveform amplitudes to be positive using the *ReLU*(·) function. In contrast, ICA source waveforms exhibit alternate polarity amplitudes at different times. Either of these behaviours may be preferred for modelling certain kinds of sources. A legitimate multi-phasic biological source waveform comes from eye blinking. As the eyelids close, the eyeballs articulate within their sockets, causing the electric dipoles between anterior chambers and posterior eyeballs to rotate back and forth. Blinking therefore generates source potential waveforms with polarity inversions due to large eyeball equivalent dipole orientation changes. This process can be represented by a single IC that allows positive and negative amplitudes. However, the RNN represents the same eye-blink with two source waveforms; one for positive and another for negative scalp amplitudes. An example of this is given in Figure S6, which was obtained by applying RNN and ICA methods to the same ERP CORE data before removing eye-blink artifacts with ICA and IClabel [28].

Eye-blinks can distort ERP waveforms when they are systematically linked to experimental events [3]. For example, if blinking tends to occur immediately after visual stimuli and/or is related to the time interval between stimuli. Source waveforms associated with each ERP in Figure S6 may thus reflect different stimulus timings and their systematic relation to eye-blinks. For instance, the active visual oddball (P3) paradigm had the longest interstimulus interval, and the corresponding eye-blink source signals had the earliest onset and largest magnitude. Furthermore, the eye-blink source waveforms for MMN had comparatively low amplitude and did not follow the same pattern as those associated with visually-evoked difference waves. The RNN also appeared to mix part of the eye-blink with P3b, indicating that the eye-blink artifact was systematically related to rare target stimuli.

Neural sources of ERP components do not exhibit dynamic changes in dipole orientation like the eye-blink because cortical tissue does not articulate. Voltage polarity at scalp electrodes due to cortical pyramidal cells is influenced by four factors: (i) orientation of the active patch of cortex relative to the scalp, (ii) whether summed post-synaptic potentials are mostly excitatory or inhibitory, (iii) whether synaptic potentials are at distal or apical dendrites, and (iv) the location of active and reference EEG electrodes [1], [3]. Factor (iv) is fixed during an experiment, and re-referencing to the montage mean post-hoc reduces sensitivity of recordings to neural activity detected by the original reference electrode. Each neural source can be assumed to have its own patch of cortex, be either inhibitory or excitatory, and act mainly on apical or distal dendrites. Under these assumptions, biphasic source waveforms imply either different patches of cortex, switching from excitatory to inhibitory activity, or moving site of activation from apical to distal dendrites. If any of these conditions arise, resulting scalp potential polarity changes could be fairly attributed to distinct neural sources, insofar as different neurophysiological mechanisms are responsible for the change in polarity. Therefore, each independent neural source can be assumed to have fixed orientation and polarity, with time-varying magnitude. Providing that dipoles do not physically articulate, such as during eye-blinking, rectified activation of RNN sources is thus an advantage in representing distinct neural sources and their corresponding scalp maps. However, this behaviour of the RNN could easily be changed by applying different activation functions to its penultimate layer outputs.

#### 4.2.3. Utility as a computational model

ICA is enacted by a single-layer artificial neural network trained to minimize shared information among its output units [30], [31]. ICA thus transforms EEG signals into ICs. The reverse operation of transforming ICs into EEG signals is made possible by inverting the weight matrix used to convert EEG into ICs. This inverse matrix is therefore equivalent to the scalp distribution of each IC [12]. The RNN has five layers of artificial neurons that perform a sequence of nonlinear transformations on input signals representing events to produce waveforms matching the grand-average of waveforms used as labels during model training [21], [22]. The output layer of the RNN learns weights to transform sources into ERP signals, which performs the same role as the ICA inverse matrix.

The ICA method does not consider temporal dependencies within the data that are crucial for RNN modelling. Extracting temporal relationships from neural data makes the RNN a more attractive model of ERP generation than the comparatively simple ICA transformation, particularly because the RNN’s computational structure is amenable to further study. For instance, recurrent weights from *SimpleRNN* 4 layer, *W*_r_^(4)^, plotted in Figure S7, reflect functional connectivity between sources and may be studied to gain insights into interactions among sources responsible for generating event-related neural signals. Moreover, it is theoretically feasible to examine shared drivers of sources from connected signals in preceding layers of the network (Figure 5), although new methods are needed for this.

### 4.3. Limitations

RNN and ICA methods incorporate stochastic optimization, thus producing variable solutions with different initial conditions [22], [30]. Source signal order is particularly variable across solutions, although time-courses and scalp maps tend to be fairly reproducible [12]. However, the computational cost of the two methods differs more considerably. Implementing ICA effectively involves training a one-layer feed-forward neural network, whereas the RNN method involves training a five-layer recurrent neural network. The computational requirements of the RNN method are therefore higher than that of implementing ICA.

For EEG artifact correction, ICA is applied to continuous EEG from a single-subject [3], [13]. One reason for this is that ICA assumes channel properties are stationary. However, group ICA may be more likely to violate this assumption of stationarity. It is also desirable to have more data points for training the neural network used in ICA to minimize mutual information among its outputs. Averaged ERP waveforms in ***Y*** provided 15,049 total time samples (from 28 EEG channels), equivalent to 2.5 min of data, which is slightly below the minimum recommended number of samples for implementing ICA (i.e., ≥ 15,680 = 20 x *N_eeg_*^2^; [12]). It could also be argued that *Extended Infomax* is not the ideal ICA algorithm for benchmarking [47]. Therefore, higher MI and SE for ICA sources than RNN sources could be partly explained by suboptimal application of one ICA algorithm to grouped ERP data.

It would be possible to estimate the position, orientation and amplitudes of each of the RNN sources by fitting equivalent current dipole models to their scalp projections [12], [48], [49]. However, arguably too few EEG electrodes were used for accurately locating sources [50]. A higher density montage would entail more ICA sources, but could also yield more RNN sources if scalp topographies of underlying neural sources become distinguishable. Nevertheless, even after being well-separated by RNN or ICA methods, more electrodes would be preferable for fitting equivalent current dipoles.

## 5. Conclusions

The RNN blind source separation method generates plausible source waveforms and scalp topographies. This technique is effective for generalizing grand-average ERP difference waveforms, but could easily be adapted for single-subject ERP data by using single-trial EEG signals as model labels. Some of the sources proposed by the RNN are very similar to ICs from ICA applied to the same data, quantified by mutual information and correlation between source waveforms and correlation between scalp maps. Despite these similarities, RNN sources were smoother, sparser, more ERP-specific, and had less ambiguity between source waveform and projected scalp potential polarities. After training the RNN, it provides a mathematical model describing transformations from representations of psychophysiological events into corresponding ERP waveforms. There is scope for extending this approach and developing analytical methods to potentially derive insights from the trained RNN about computational processes underlying event-related neural processing.

## Acknowledgements

JAO and EWL would like to thank Emily S. Kappenman and Steven J. Luck for providing training at the 2023 UC-Davis/SDSU ERP Boot Camp funded by the U.S. National Institute of Mental Health, and all of the authors thank them for making the data used in this study freely available. Research reported in this publication was supported by the National Institutes of Health under Award Number NIAMS K23AR083171 (EWL). This work was financially supported by King Mongkut’s Institute of Technology Ladkrabang [2567-02-01-061].

## Data availability

Data and code from this study will be shared publicly via an Open Science Foundation repository.

## Author contributions (CRediT statement)

JAO: Conceptualization, Methodology, Software, Validation, Investigation, Data Curation, Writing - Original Draft, Writing - Review & Editing, Visualization, Supervision HAS, NA, PK, PC, and TK: Software, Validation, Investigation, Data Curation EWL: Investigation, Data Curation, Writing - Review & Editing, Visualization

## Supplementary Information

**Table S1.**
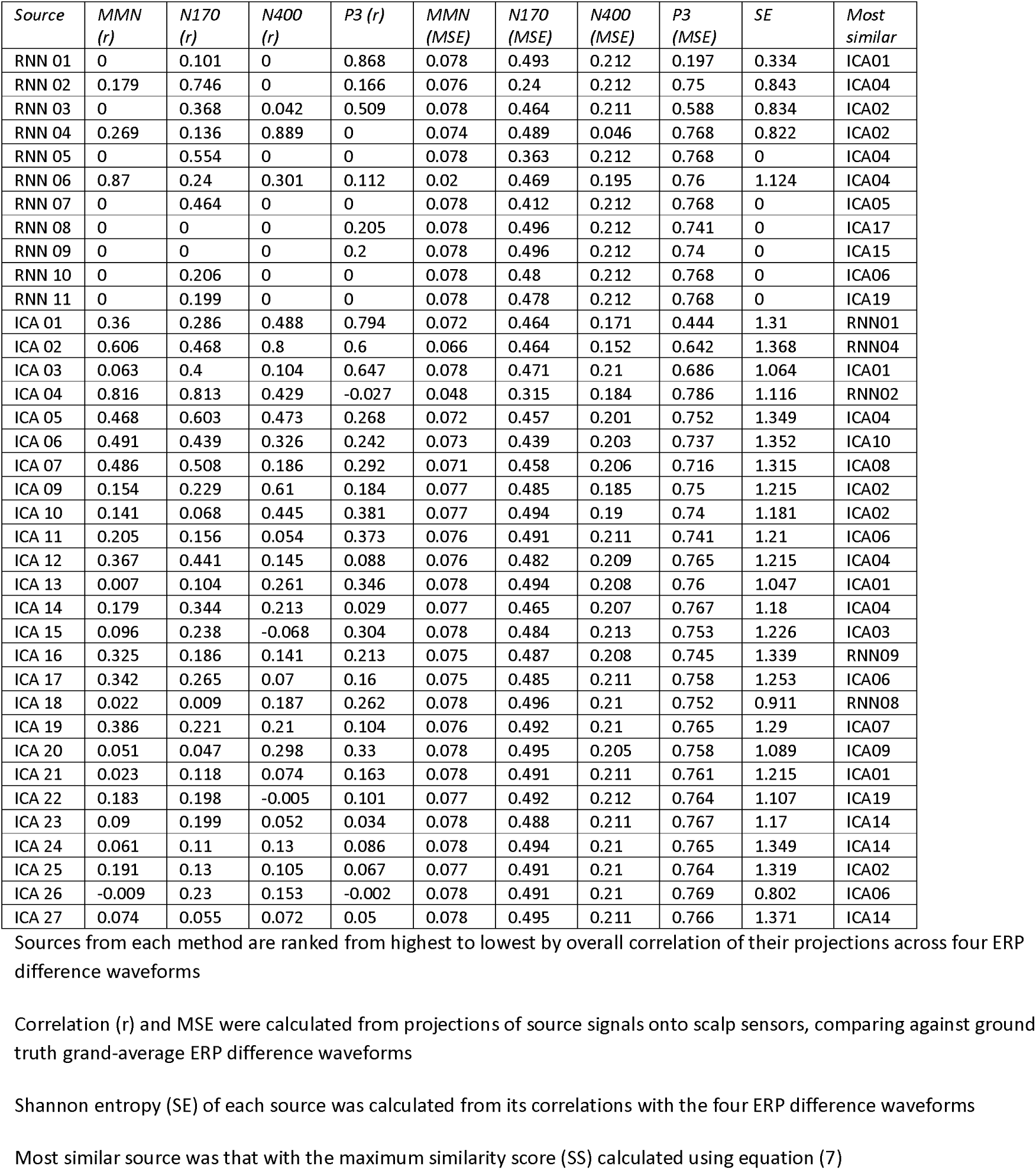
Analysis of RNN and ICA sources.

**Figure S1.**
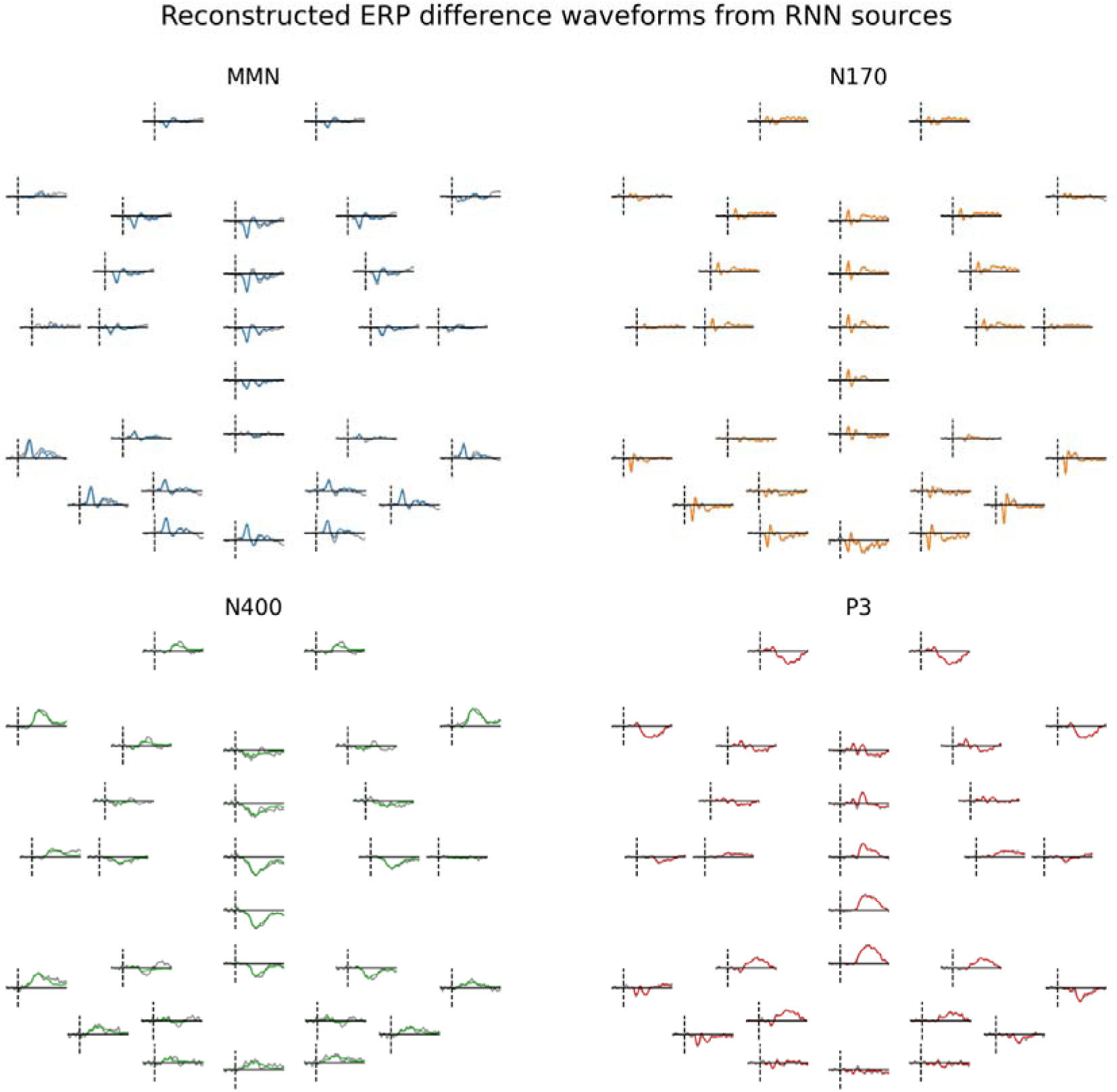
Reconstructed difference waveforms from RNN sources plotted at 28 EEG channels. Reconstructed waveforms are plotted in colour and ground-truth waveforms are plotted in black, although two traces on most plots are difficult to distinguish because they are overlapping. All of the electrodes in each quadrant are plotted with the same y-axis scale.

**Figure S2.**
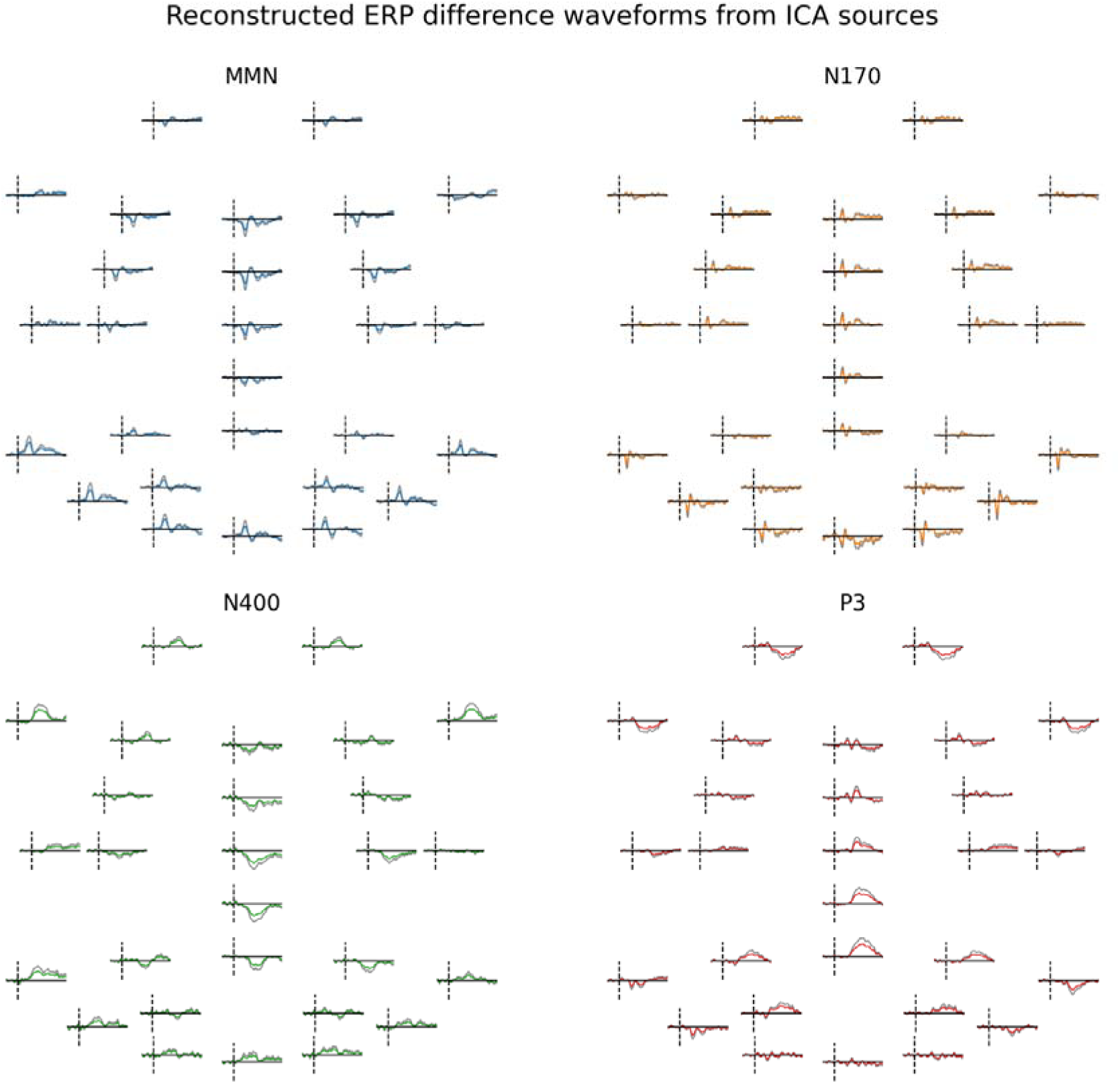
Reconstructed difference waveforms from ICA sources plotted at 28 EEG channels. Reconstructed waveforms are plotted in colour and ground-truth waveforms are plotted in black. Y-axis scales are the same for each electrode in the montage, but different for different ERP difference waveforms (MMN, N170, N400, and P3). These reconstructed waveforms have higher MSE compared with those from RNN sources, but otherwise are comparably well-correlated with ground-truth waveforms.

**Figure S3.**
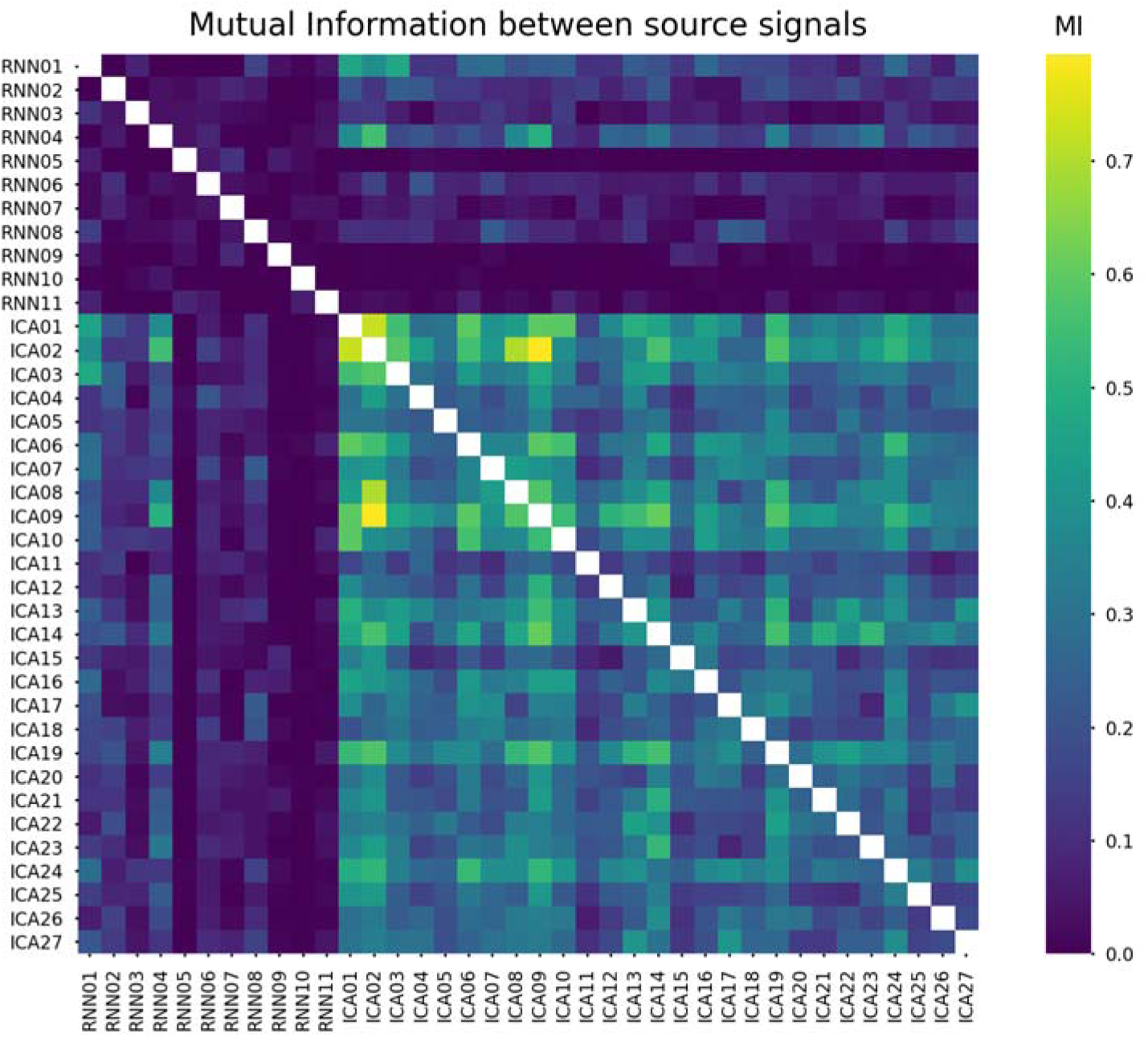
Pairwise mutual information between RNN and ICA sources. MI is generally higher between ICA sources than it is for RNN sources. There also tends to be higher MI between RNN and ICA sources than within RNN sources. These dependencies are reflected by signal waveforms for these sources.

**Figure S4.**
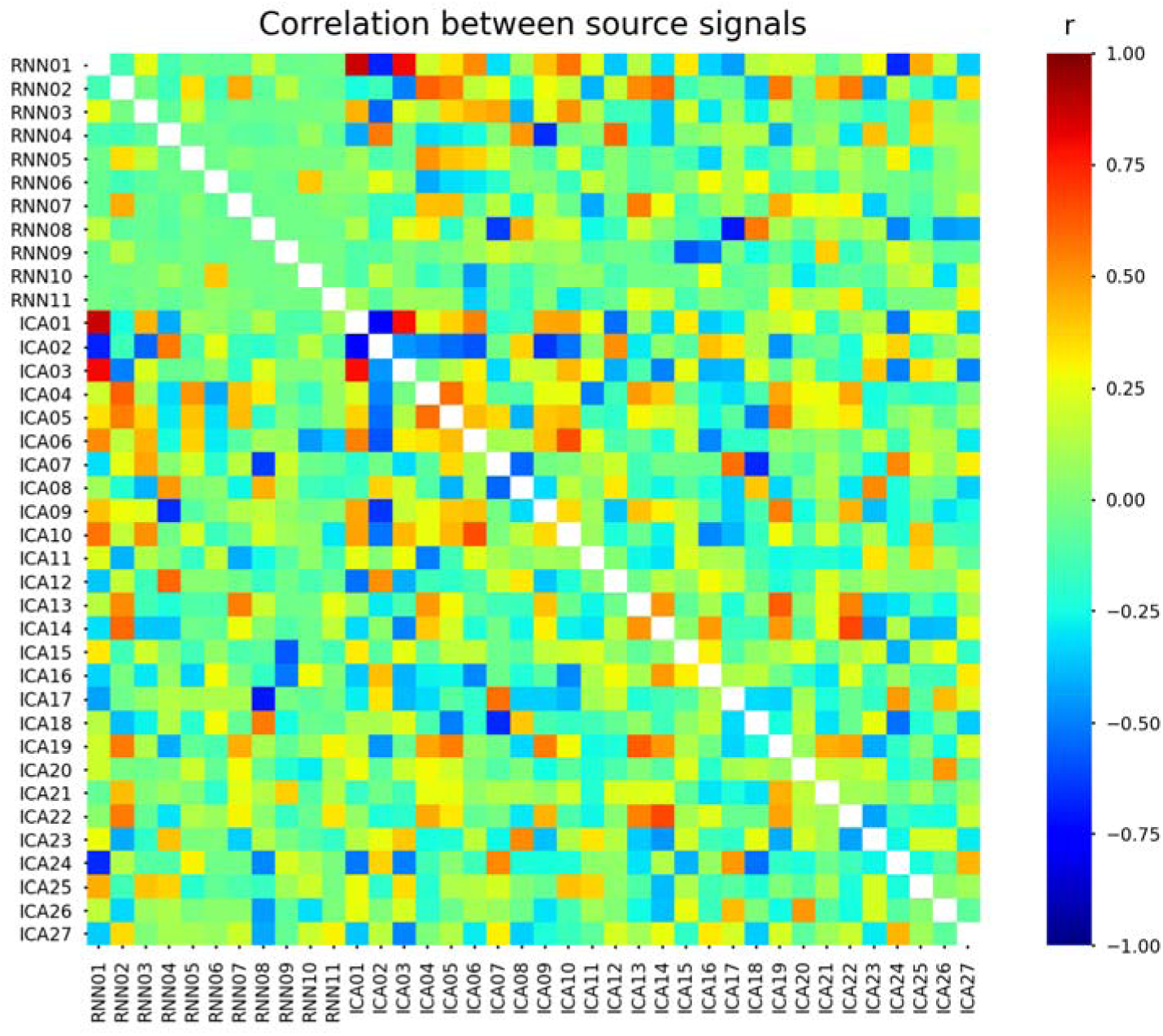
Pairwise correlations between signals for each RNN and ICA source. A similar pattern seen from MI analysis is observed for correlation between source signals. However, signals can be anticorrelated because ICA allows positive and negative source amplitudes, which may reflect inversions of RNN sources. Correlations within ICA sources are higher than those within RNN sources, and RNN sources tend to have greater correlations with ICA sources than with RNN sources.

**Figure S5.**
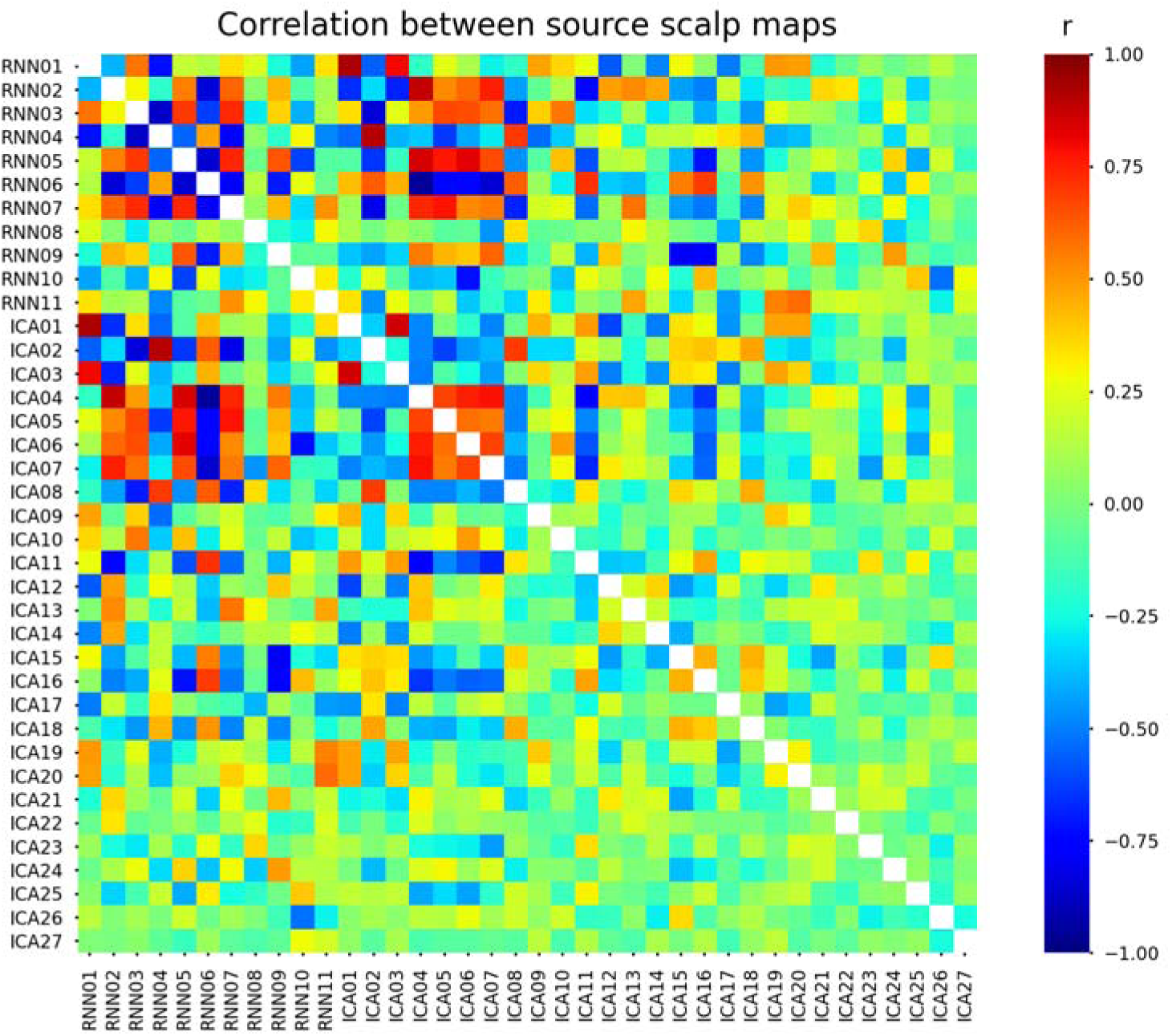
Pairwise correlations between scalp maps for each RNN and ICA source. There are high correlations within and between RNN and ICA sources, reflecting similarity of scalp distributions. For RNN sources, negative correlations indicate opposite polarity contributions to scalp potentials. However, for ICA sources this relationship is ambiguous because ICA source signals can have biphasic polarity.

**Figure S6.**
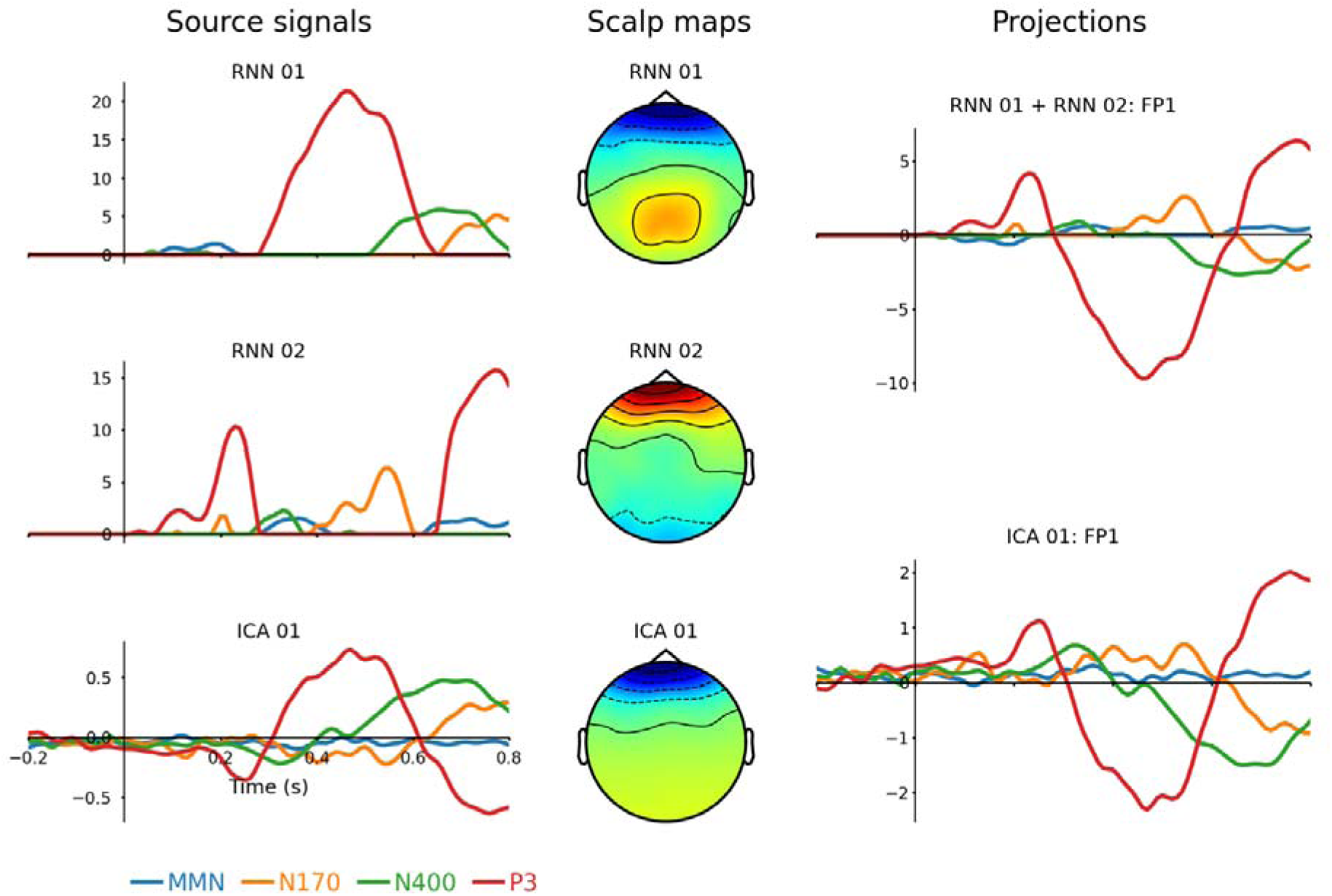
Example of an eye-blink source represented by RNN and ICA methods. This analysis was performed on data before correcting eye-blink artifacts. Source ICA 01 captures both phases of activity caused by changes in electric dipole orientation as the eye articulates in its socket while blinking. RNN 01 captures negative portions and RNN 02 captures positive portions of the blink artifact; positive and negative polarities given in terms of scalp projections at electrode FP1. The summed projections from RNN 01 and RNN 02 are highly correlated with the projection from ICA 01. Source waveforms and projections are plotted on the same time range. Differences in the latency of this systematic eye-blink artifact for N170, N400 and P3 may reflect differences in stimulus duration and interstimulus-interval (ISI) in each of these paradigms (i.e., N170: 0.3 s duration, 1.1-1.3 s ISI; N400: 0.2 s duration, 0.9-1.1 s ISI before target words; P3: 0.2 s duration, 1.2-1.4 s ISI).

**Figure S7.**
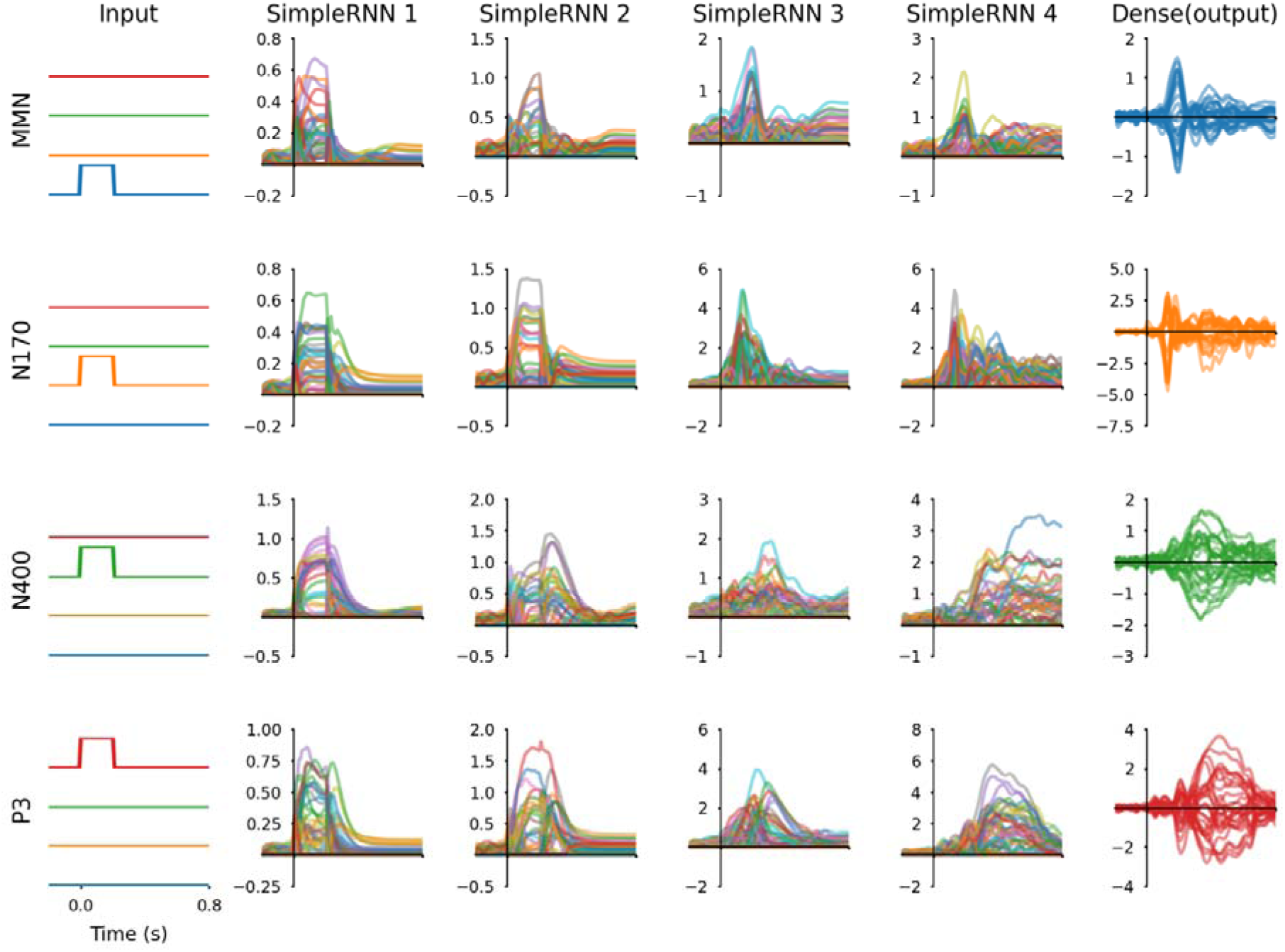
Inputs, hidden unit activations, and output signals from the RNN after training phase 1 (without L1 regularization applied to *SimpleRNN* 4 layer). All 64 hidden units from SimpleRNN 4 are involved in producing outputs that match ERP difference waveforms; this makes it possible to use the RNN method to separate more sources than the number of EEG channels. All waveforms are plotted from −0.2 s to 0.8 s about stimuli onsets that occurred at 0.0 s. Hidden unit activation waveforms have arbitrary units and output waveforms have microvolt units.

**Figure S8.**
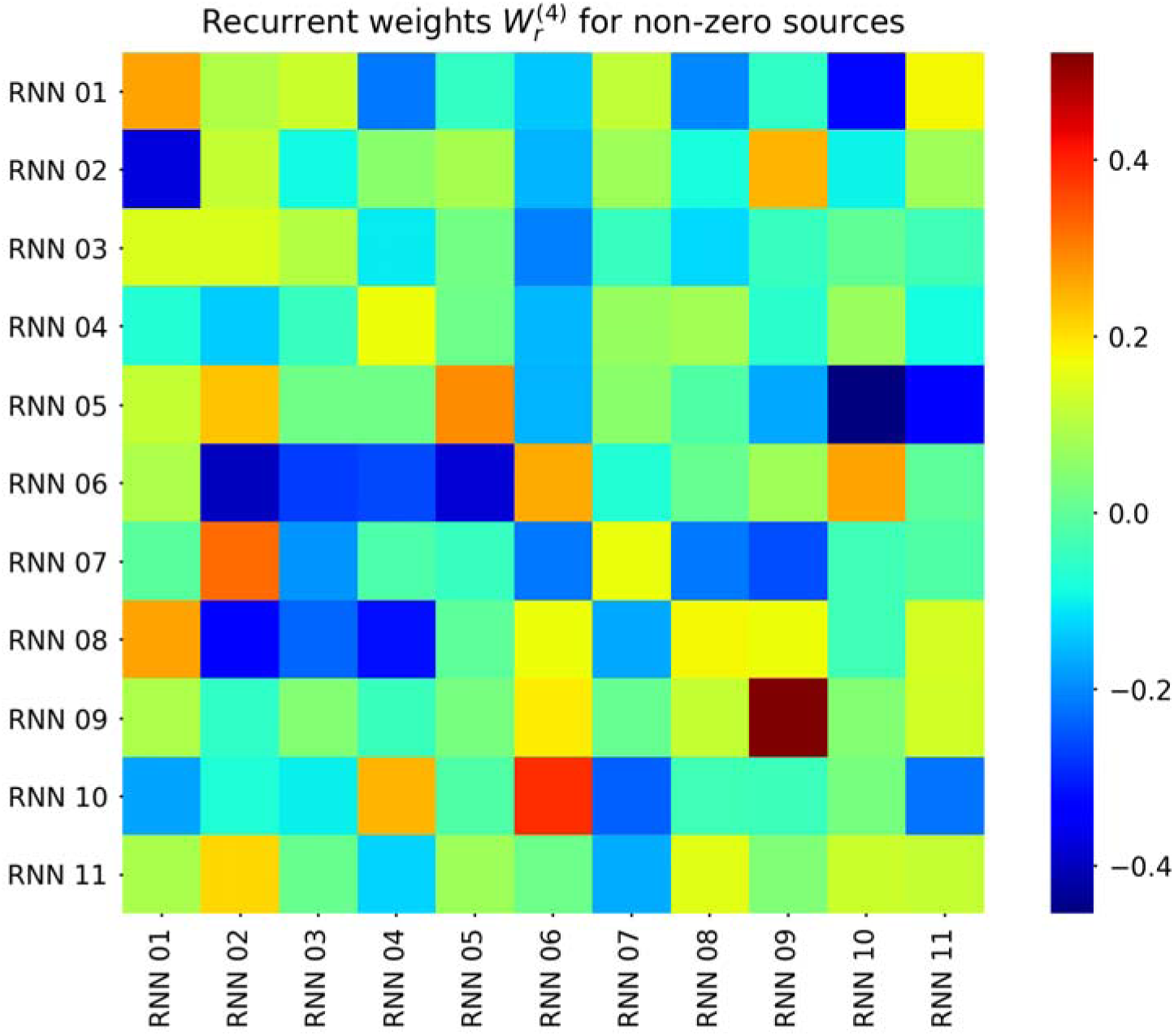
Recurrent weights from the *SimpleRNN 4* layer of the RNN. These recurrent weights determine the influence of *affector* source at time in each row with *affected* source at time in each column; hence the direction of influence is from row to column, as presented in this figure. Negative values indicate the tendency for an affector source to diminish or suppress activity of the affected source, while positive values indicate a tendency for an effector source to enhance or induce activity of the affected source. These recurrent weights thus reflect functional connectivity among sources. It is important to remember that inputs from the preceding hidden layer (*SimpleRNN* 3) also influence the activity of these sources..

